# Mining Large Heterogeneous Cancer Data Sets Using Boolean Implications

**DOI:** 10.1101/045021

**Authors:** Subarna Sinha, David L. Dill

## Abstract

Boolean implications (if-then rules) provide a conceptually simple, uniform and highly scalable way to find associations between pairs of random variables. In this paper, we describe their usage in mining associations from large, heterogeneous cancer data sets. Next, we illustrate how Boolean implications were used to discover a new causal association between a mutation and aberrant DNA hypermethylation in acute myeloid leukemia as well as the therapeutic implications of this discovery. We conclude with a brief description of how Boolean implications can be extracted from a given data set.

## I. MINING ASSOCIATIONS IN LARGE CANCER DATA SETS

Large-scale cancer genome projects including The Cancer Genome Atlas (TCGA) (http://cancergenome.nih.gov/) are generating an unprecedented amount of multidimensional data using high-resolution microarray and next-generation sequencing platforms. There are opportunities for mining these data sets that can yield insights that would not be apparent from smaller, less diverse data sets. Obtaining the full value of these data requires the ability to find associations between heterogeneous data types.

In this paper, we describe the use of Boolean implications [1] to find pairwise associations in heterogeneous cancer data sets. Boolean implications are if-then rules. The distribution of points in a scatterplot of two variables in a Boolean implication is L-shaped instead of linear (Fig 1). There are four Boolean implications: (1) if A is low, then B is low (*LOLO*), (2) if A is high, then B is low (*HILO*), (3) if A is low, then B is high (*LOHI*), (4) if A is high, then B is high (*HIHI*). Boolean implications can also be interpreted according to set theory. The *HIHI* Boolean implication “if A is high, then B is high” means that “the set of samples where A is high is a subset of the set of samples where B is high”. The *HILO* implication “if A is high, then B is low” means that “the set of samples where A is high is mutually exclusive with the set of samples where B is high”. Previous work [1] showed that a large number of Boolean implications are present in gene expression data. Previous applications of Boolean implications in biology were mainly to understand development [2, 3].

**Figure 1.**
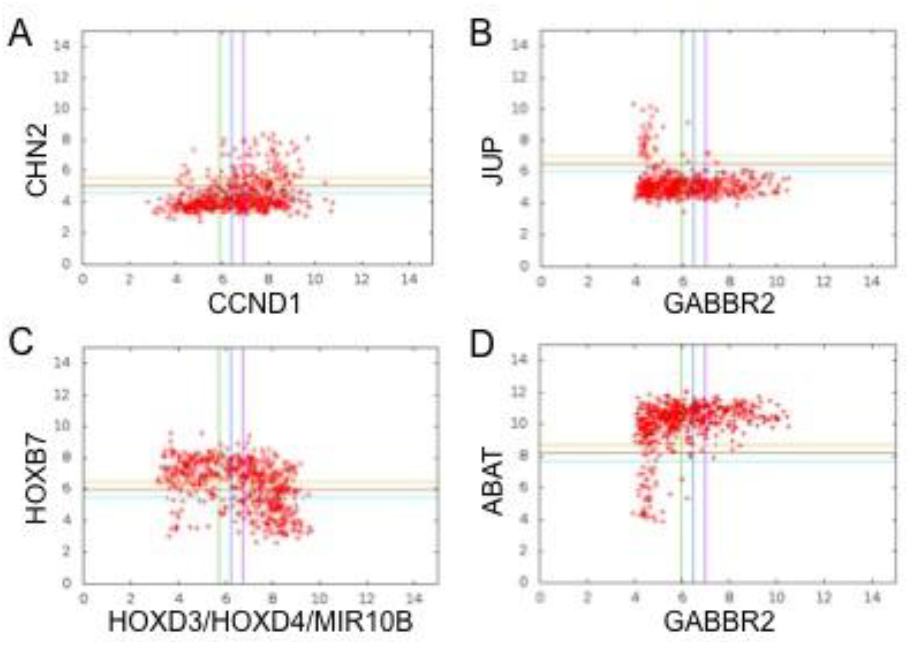
Boolean Implications illustrated using data from gene expression arrays. (A) LOLO (if CCND1 is low, then CHN2 is high) (B) HILO (if GABBR2 is high, then JUP is low) (C) LOHI (if HOXD3 is low, then HOXB7 is high) (D) HIHI (if GABBR2 is high, then ABAT is high)

We hypothesized that Boolean implications would be useful in the context of mining heterogeneous cancer data sets because (1) they can expose subset and mutual exclusion relationships, both of which have L-shaped scatterplots between related variable pairs; and (2) they can expose relationships between variables from different assays. Accordingly, we extracted Boolean implications between mutation, copy number alteration, DNA methylation and gene expression for several large TCGA data sets. Our experiments show that large numbers of implications exist. Experimental comparisons with existing commonly used methods to identify pairwise associations in biological data such as the *t* test, correlation and Fisher’s exact test revealed that many Boolean implications are missed by other methods [4]. Furthermore, many of the relationships found by Boolean implications captures key aspects of cancer biology.

## II. APPLICATION OF BOOLEAN IMPLICATIONS TO DERIVE NEW INSIGHTS IN CANCER BIOLOGY

Boolean implications can be used to derive insights for a variety of problems in cancer biology. Boolean implications from the TCGA data have revealed *cis* relationships between copy number alteration, DNA methylation and expression of genes, a new hierarchy of mutations and recurrent copy number alterations, loss-of-heterozygosity of well-known tumor suppressors, and the association between mutations and aberrant DNA hypermethylation in cancer. Below, we describe the application of Boolean implications to infer mutation-specific DNA methylation signatures.

Aberrant changes in DNA methylation are known to play a major role in the evolution of multiple cancers, but the molecular events responsible for perturbing methylated genomic landscapes have not been completely characterized. In order to identify somatic mutations in cancer that are directly linked to DNA CpG methylation, we developed a Boolean-implication based algorithm to systematically analyze the TCGA mutation and DNA methylation data (Fig 2A). As a first step, we applied the algorithm to the 16 recurrent mutations in acute myeloid leukemia (AML), which is known to have several molecular drivers of DNA methylation. Consistent with previous findings [4, 5], the algorithm identified relationships between mutations in *IDH2* and *CEBPA* and DNA hypermethylation in AML. Interestingly, we found that the *WT1* mutation was associated with a predominance of *HIHI* Boolean implications with DNA methylation, suggesting a new association between mutation in the Wilms’ Tumor 1 (WT1mut) gene and CpG hypermethylation (Fig 2B). Introduction of WT1mut into wildtype THP1 AML cells induced DNA hypermethylation in the same set of genes, confirming WT1mut to be causally associated with DNA hypermethylation in AML (Fig 2C). Methylated genes in WT1mut AML cells were highly enriched for polycomb repressor complex 2 (PRC2) targets, which were also aberrantly repressed in WT1mut primary patient samples.

**Figure 2.**
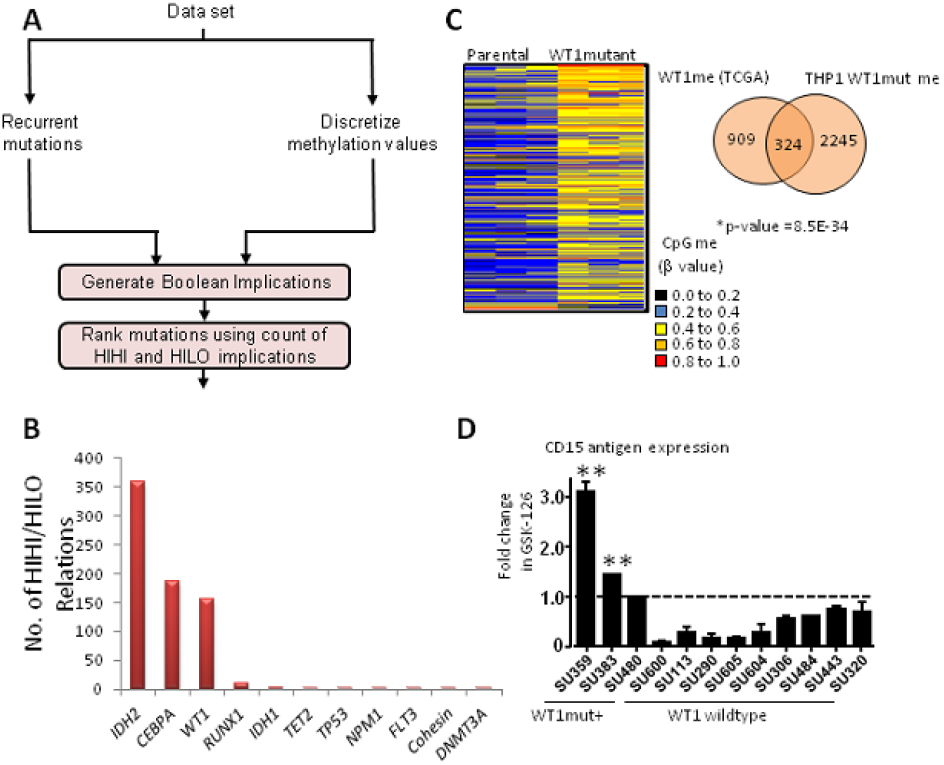
Mutant WT1 is Causally Associated with DNA Hypermethylation in Acute Myeloid Leukemia (A) Boolean Implication-Based Analysis of Somatic Mutation and DNA Methylation Data (B) WT1 mutation (C) Introduction of WT1 mutation in THP1 cells induces increase in methylation; Statistically significant overlap exists between genes methylated in WT1mut TCGA samples and WT1mut THP1 cells. (D) Summary of fold increase in CD15 antigen expression after GSK-126 treatment in vitro compared to DMSO for WT1mut+ AMLs and nine WT1 wildtype AMLs.

Furthermore, treatment of primary WT1mut AML cells with PRC2/EZH2 inhibitors (GSK-126) promoted myeloid differentiation, suggesting EZH2 inhibitors may be active in this AML subtype (Fig 2D). More details on this work can be found in [6]. The Boolean implication-based analysis is generalizable and can be applied to analyze mutation-specific DNA methylation signatures in other cancers as well. Application of the method to other TCGA cancers-bladder, breast, head and neck squamous, renal clear cell, lower grade glioma, lung adenocarcinoma, lung squamous, ovarian and uterine - revealed new associations between somatic mutations and aberrant DNA hypomethylation including (i) *STAG2* (a member of the cohesin complex) in bladder cancer and AML, (ii) *DNMT3B* (an alternative de-novo DNA methyltransferase) in lung adenocarcinomas and (iii) *KDM5C* (a histone demethylase) clear cell cancer.

In summary, Boolean implication-based analysis can be used to identify mutation-specific DNA methylation signatures in cancer. Furthermore, we demonstrate that deciphering mutation-specific methylation patterns can lead to therapeutic insights

## III. QUICK GUIDE TO EXTRACTING BOOLEAN IMPLICATIONS

The first step in the extraction of Boolean implications is to convert all attributes to Boolean variables. In our analysis, gene expression and DNA methylation data were discretized using StepMiner [7]. Subsequently, Boolean implications were detected using a statistical test consisting of two parts: (1) an independence test (Fisher’s exact test) to detect nonrandom associations, (2) then the sparsity test checked for sparseness of a specific quadrant using a maximum-likelihood estimate of the error rate for the points in the sparse quadrant [1]. An implication was considered significant if the first statistic was greater than a cutoff threshold and the error rate was less than 0.1. Note that the sparsity test (step 2) distinguishes a Boolean implication from simple non-independence of variables.

Given the large number of attributes and even larger number of potential relationships, it was necessary to evaluate the significance of the relationships discovered by the above algorithm. An estimate of the false discovery rate (FDR) was obtained by randomly permuting the values for each attribute independently, and then extracting the Boolean implications as above. This analysis was repeated 50 times to compute the average number of Boolean implications in the randomized data. The FDR was the ratio of the average number of Boolean implications in the randomized data and the original data. The cutoff thresholds for the Boolean implication test were set to obtain an acceptable FDR.

### B. Type of settings in which these methods are useful

Boolean implications can be used to extract pairwise relationships, in particular subset and mutual exclusion relationships, in any large heterogeneous data.

## References

[1] D. Sahoo, DL Dill, AJ Gentles, R Tibshirani and S Plevrits, “Boolean implication networks derived from large scale, whole genome microarray datasets”, in Genome Biology, 9(10):R157, 2008.

[2] D. Sahoo, J. Seita, D. Bhattacharya, M. Inlay, I. Weissman and DL. Dill, “MiDReG: a method of mining developmentally regulated genes using boolean implications”, in Proceedings of the National Academy of Sciences 107(13): 5732-5737, 2010.

[3] P. Dalerba, T. Kalisky, D. Sahoo, P. Rajendran, ME. Rothenberg et al, “Single-cell dissection of transcriptional heterogeneity in human colon tumors”, in Nature Biotechnology, 29(12): 1120-1127, 2011.

[4] ME. Figueroa, O. Abdel-Wahab o, C. Lu, et al., “Leukemic idh1 and idh2 mutations result in a hypermethylation phenotype, disrupt tet2 function, and impair hematopoietic differentiation”, in Cancer Cell, 18(6):553–567, 2010.

[5] ME. Figueroa, S. Lugthart, Y. Li, et al., “DNA methylation signatures identify biologically distinct subtypes in acute myeloid leukemia”, in Cancer Cell, 17(1):13–27, 2010.

[6] S. Sinha, EK. Tsang, H. Yao, M. Meister and DL. Dill, “Mining TCGA Data Using Boolean Implications”, in PLoS One, 9(7):e102119, 2014.

[7] S. Sinha, D. Thomas, L. Yu, AJ. Gentles, N. Jung, MR. Corces-Zimmerman, SM. Chan, A. Reinsich, AP. Feinberg, DL. Dill and R. Majeti, “Mutant WT1 is associated with DNA hypermethylation of PRC2 targets in AML and responds to EZH2 Inhibition”, in Blood, 125(2):316–26, 2015.

[8] D. Sahoo, DL. Dill, R. Tibshirani and S. Plevritis, “Extracting binary signals from microarray timecourse data”, in Nucleic Acids Research, 35(11): 3705-372, 2007.

